# Circadian Coupling Orchestrates Cell Growth

**DOI:** 10.1101/2024.05.18.594797

**Authors:** Nica Gutu, Malthe S. Nordentoft, Marlena Kuhn, Carolin Ector, Anna-Marie Finger, Mathias Spliid Heltberg, Mogens Høgh Jensen, Ulrich Keilholz, Achim Kramer, Hanspeter Herzel, Adrián E. Granada

**Affiliations:** Charité Universitätsmedizin Berlin, Berlin, Germany; Humboldt Universität zu Berlin, Berlin, Germany; Niels Bohr Institute, University of Copenhagen, Copenhagen, Denmark; German Cancer Consortium (DKTK), Berlin, Germany; Institute for Theoretical Biology, Berlin, Germany

**Keywords:** circadian clock, cell cycle, cell growth, biological oscillators

## Abstract

Single-cell circadian oscillators exchange extracellular information to sustain coherent circadian rhythms at the tissue level. Within cells, the circadian clock and the cell cycle couple, yet the mechanisms governing this interplay remain poorly elucidated. Here, we study the role of extracellular circadian communication in the intracellular coordination between the circadian clock and the cell cycle. We demonstrate that the loss of extracellular circadian synchronization disrupts circadian and cell cycle coordination within individual cells, impeding collective tissue growth. Coherent circadian rhythms yield oscillatory growth patterns, unveiling a global timing regulator of tissue dynamics. Knocking down core circadian elements abolishes observed effects, highlighting the central role of circadian clock regulation. Our research underscores the significance of tissue-level circadian disruption in regulating proliferation, thereby linking disrupted circadian clocks with oncogenic processes. These findings illuminate the intricate interplay between circadian rhythms, cellular signaling, and tissue physiology, enhancing our understanding of tissue homeostasis and growth regulation in both health and disease contexts.

## Introduction

Due to Earth’s rotation, organisms experience cyclical environmental changes associated with day and night cycles. In response, most organisms have evolved an internal circadian clock to anticipate and coordinate physiological and metabolic processes with the environmental cycles^1,2^. In humans, the circadian clock operates through a hierarchical organization, wherein a master clock situated in the hypothalamic suprachiasmatic nucleus coordinates the activities of peripheral autonomous clocks distributed across the body^3^. At the single-cell level, the molecular clock machinery relies on auto-regulatory transcription-translation feedback loops that consist of both positive and negative elements, driving the cyclic expression of clock genes and proteins with a period of around 24 hours^4,5^. Coupling between single-cell circadian oscillators results in collective tissue-level rhythms with tissue-specific oscillatory strength and entrainment properties^6,7^. The interaction between single-cell oscillators within peripheral tissues involves intercellular coupling mechanisms facilitated by paracrine communication pathways, where the transforming growth factor–β (TGF-β) acts as the intermediary transmitting paracrine synchronization signals to the molecular clock machinery^8^. Thereby, these cell-autonomous single-cell circadian oscillators can exchange information and couple to secure synchronization of the cellular network and avoid the disruption of circadian tissue functions (Fig. 1).

**Figure 1:**
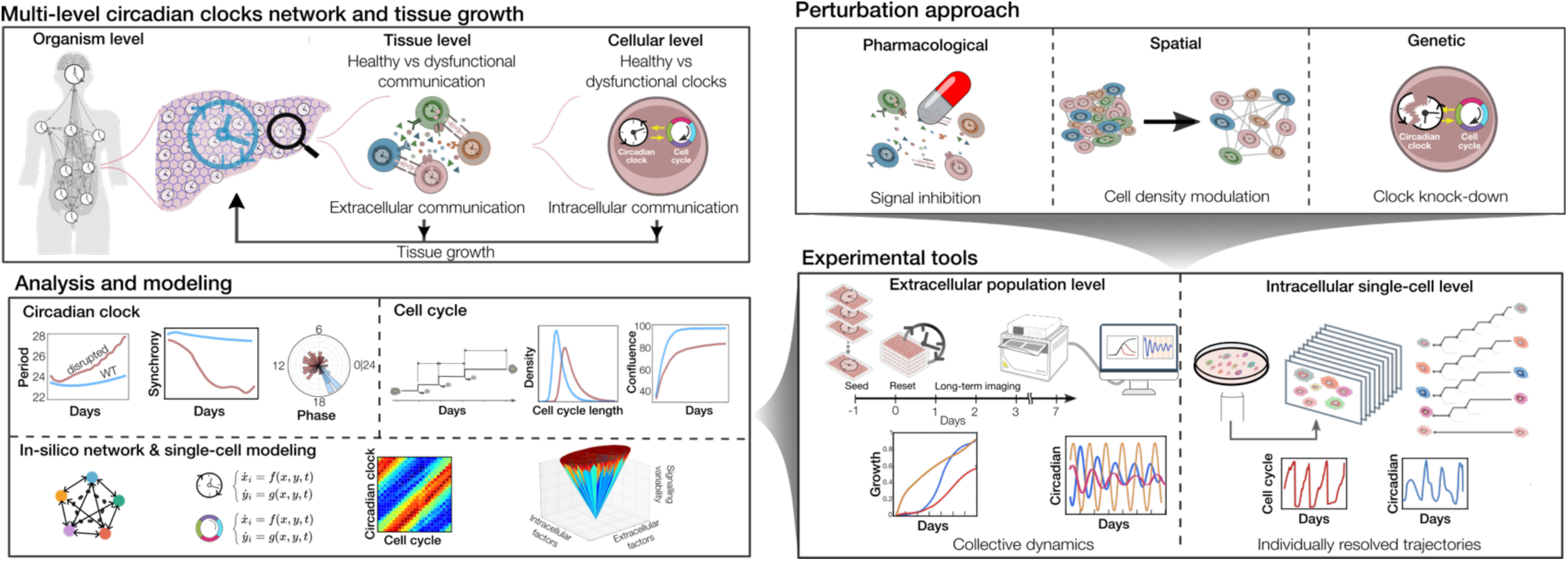
Framework to identify the interplay between extracellular and intracellular communication. Top left: sketch of the multi-level networks of the circadian clocks within a human body from the organism to the single-cell level. The master clock coordinates the peripheral autonomous clocks distributed across every tissue in the body. At the tissue level, the peripheral cell-autonomous clocks communicate with each other and constitute networks of circadian clocks. At the single-cell level, the circadian clock and the cell cycle communicate. Top right: sketch of the perturbation approaches used in this study were: pharmacological to inhibit the circadian signal exchanged among cells, spatial to modulate cell density to reduce the amount of circadian signal present in the media, and genetic to knock down the circadian clock. Bottom right: sketch of the long-term live imaging at population and single-cell levels experiments used to study the collective dynamics and individual resolved trajectories of the circadian clock and the cell cycle. Bottom left: sketch of the metrics from signaling dynamics used to analyze the collected data and build a mathematical model.

In proliferating cells, another main regulatory biological process unfolds, the cell cycle, with a comparable characteristic period of 24 hours. This process involves a series of ordered and regulated events involving cell growth, DNA replication, and division, encompassing distinct phases such as G1 (gap 1), S (synthesis), G2 (gap 2), and M (mitosis)^9–12^. The stepwise progression through recurring phases of activation and repression and successive divisions can be conceptualized as another periodic process^13^.

Hence, at the single-cell level, two periodical biological processes with analogous periods coexist and engage in mutual influence^14–19^, resembling two oscillators capable of coupling and synchronization leading to important physiological implications in tissue homeostasis or cancer (reviewed in^20^). Comprehensive studies across diverse systems sought to explore these biological oscillators and unravel the mechanisms underlying their interactions. However, numerous questions persist including the nature and directionality of the coupling between the circadian clock and the cell cycle and, most importantly how cellular factors regulate this relationship across tissues.

This study investigates how extracellular circadian clock synchronization plays in the intracellular communication between the cell cycle and the circadian clock. To address this question, we built a mathematical model and performed long-term live-imaging recordings at the single-cell and population level of the human osteosarcoma cell line (U2OS), a well-established in vitro model of human peripheral clocks. We employed various strategies to disrupt extracellular circadian coupling with pharmacological perturbations, cell density control to diminish the amount of exchanged circadian information in the media, and genetic disruption of the circadian clock (Fig. 1).

Our results showed that the population’s circadian and cell cycle coordination can be destroyed by diminishing the circadian extracellular synchronization using a TGF-β secretory pathway inhibitor or through the seeding density. Moreover, our findings indicated that cell growth can be obstructed by disturbing extracellular circadian communication. Knockdown of fundamental circadian components led to negligible growth changes upon disruption of extracellular circadian synchronization. Finally, we observed synchronized cell division events occurring at a 24-hour periodicity within a population of well-functioning and coupled circadian clocks. Altogether, these results reveal the timing mechanism of the circadian clock as an orchestrator of tissue dynamics growth, determining the ideal time window for the cell cycle lengths to neither accelerate nor decelerate due to the 1:1 locking between the circadian and cell cycle rhythms.

## Results

### Circadian extracellular coupling enhances intracellular coupling with the cell cycle

To comprehend how the heterogeneity within an ensemble of circadian clocks influences their synchronization, we simulated a network of Poincaré oscillators^21,22^. In this ensemble, each circadian clock has an individual period and exerts an influence on all the other ones (mean-field^23^). We constructed a diagram to determine the regions of entrainment by measuring the network synchronization index while varying the dispersions of the period distribution and the coupling strengths (Fig. 2A). Here, traversing vertically this Arnold-like tongue, we can envision an increase in synchronization. For scenarios 1-3, we observed gradually damping oscillations with a period prolongation proportional to the extracellular coupling values (Figs. 2A-B). To quantify the damping of oscillations, we calculated the phase coherence which decays proportionally to the coupling strength due to an increase in the dispersion of phases over time, consistent with the observations outlined in ^8^ (Fig. 2B).

**Figure 2:**
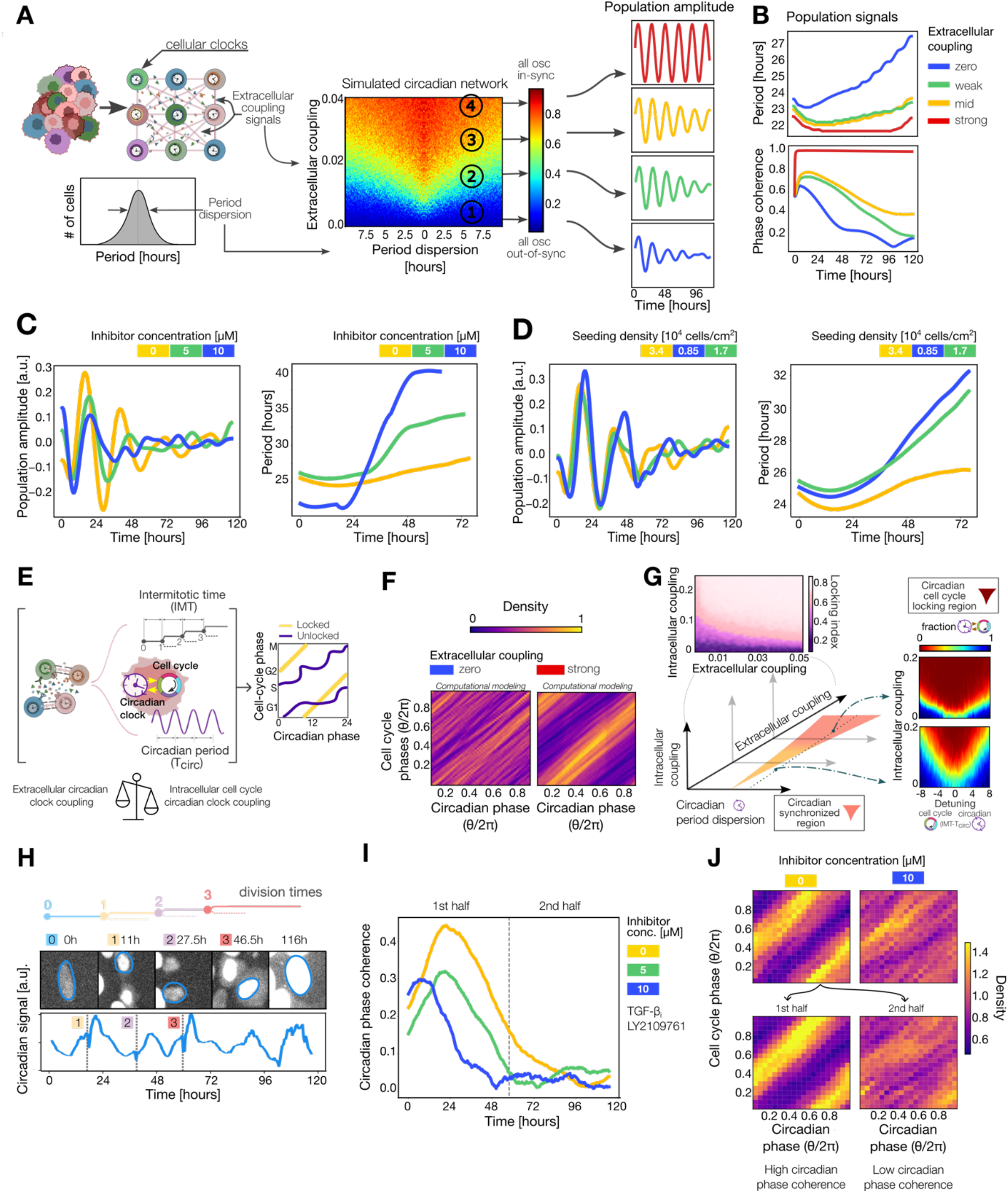
Circadian extracellular coupling enhances intracellular communication with the cell cycle. **(A)** Sketch of a network of coupled oscillators and the distribution of their periods (left). Simulation illustrating the entrainment regions of a network of circadian oscillators (right). The color coding indicates the synchronization index ranging from zero to one. For several coupling strengths, we have the corresponding simulated population amplitude after an initial resetting with a Gaussian pulse. **(B)** Period of the population circadian amplitude and the phase coherence of the simulated network corresponding to Fig. 2A after an initial resetting. **(C)** Left: experimental population circadian amplitude for different concentrations of the extracellular coupling inhibitor LY2109761. Right: period of the population circadian amplitude for different concentrations of the inhibitor. The color coding corresponds to the inhibitor concentration. Sample sizes for 0 μM (n=923), 5 μM (n=1856), and 10 μM (n=759). **(D)** Left: experimental population circadian amplitude for different seeding densities. Right: period of the population circadian signal for different seeding densities. The color coding indicates the seeding density: yellow - 34000 cells/cm^2^, green - 17000 cells/cm^2^, and blue - 8500 cells/cm^2^. The respective sample sizes are n = 923, 294, and 89. **(E)** Sketch of the interplay between the extracellular circadian coupling and the intracellular circadian clock and cell cycle communication with the corresponding periods and phase space representations with their locking state. **(F)** Simulations of the phase space representation of the circadian and cell cycle rhythms corresponding to a network of uncoupled and coupled circadian oscillators. **(G)** Three-dimensional diagram of the entrainment regions of the network of circadian oscillators intracellularly coupled to the cell cycle. Top left: simulated synchronization regions for different intracellular and extracellular coupling strengths. The color coding indicates the mean phase coherence of the individual circadian and cell cycle phase differences defined as the locking index, increasing from purple to pink. Right plots: simulation of the population intracellular entrainment region between the circadian clock and the cell cycle for a network of high and low extracellularly coupled circadian clocks. The color coding represents the fraction of locked circadian and cell cycle oscillators. **(H)** Snapshots of an untreated cell expressing the nuclear fluorescent signal corresponding to the circadian clock at the beginning, after mitosis events, and at the end time point (top). The corresponding circadian signal and division times (vertical dashed lines) are depicted below. **(I)** Experimental circadian phase coherence for different concentrations of the inhibitor. The dashed vertical line splits the experiment into two identical halves. Sample sizes are the same as in Fig. 2C. **(J)** Top: phase space two-dimensional histograms of the experimental circadian and cell cycle rhythms for two different conditions. Same sample sizes as in Fig. 2C. Bottom: phase space representation of the first (left) and the second half (right) of the experiment for the untreated cells corresponding to high and low circadian phase coherence (see Fig. S2F).

To test these predictions, we disrupted the extracellular circadian coupling with an inhibitor of the TGF-β pathway (LY2109761) or by reducing the seeding density. The circadian and cell cycle signals of the cells were consistently monitored over 5 days. Subsequently, an automated pipeline developed for ilastik^24^ was employed to track thousands of cells. Following the exclusion of noisy, non-oscillating cells (refer to Methods), around 3,000 cells remained, from which approximately 90% were oscillating with similar periods over the different conditions (Fig. S1A-D). The inhibitor of the TGF-β pathway (LY2109761) diminished the population circadian amplitude and accelerated the damping, exhibiting a characteristic period prolongation in a dose-dependent manner (Fig. 2C). Upon comparison with simulations from Figures 2A and B, we observed a parallelism between the addition of the inhibitor and the reduction of extracellular circadian coupling strength. An alternative method to attenuate coupling strength within a network of circadian clocks, interconnected via a mean-field, involves decreasing seeding density which yielded analogous density-dependent outcomes (Figs. 2D and S1E).

Next, to explore the interplay between the extracellular circadian coupling and the intracellular circadian cell cycle locking, we expanded our simulated cell network by incorporating a cell cycle oscillator coupled to each circadian oscillator (Fig. 2E). To analyze the dynamics between the circadian clock and the cell cycle, we constructed phase space representation for uncoupled and well-coupled circadian clocks (Figs. 2F and S2A and B). In an ensemble of uncoupled circadian clocks, the phases of the two rhythms are dispersed across the entire phase space, forming trajectories with different phase differences (Fig. 2F). Increasing the extracellular coupling allows the emergence of attractors indicating the conservation of a specific phase difference across the network modulated by the communication among circadian clocks (Fig. 2F).

To assess the level of intracellular locking between the circadian clock and cell cycle, we calculated the phase differences between these two rhythms for each simulated cell unit. To further quantify the degree of intracellular locking synchronization, we measured the phase coherence of these phase differences. Following the initial transient stage, the phase coherence stabilizes around a mean value depending on the interplay between the extracellular and the intracellular couplings. Here, by increasing the extracellular coupling, we can obtain higher synchronization of the circadian clock and the cell cycle communication (Figs. 2G and S2C).

In parallel, we measured the fraction of intracellular circadian and cell cycle locked cells by simulating the entrainment regions for uncoupled and well-coupled circadian clocks (Fig. 2C). Our findings revealed an enhancement in the synchronization of intracellular circadian and cell cycle communication by increasing the extracellular circadian coupling.

The single-cell level experiments perturbing the circadian extracellular coupling through the LY2109761 inhibitor, or the seeding density allowed us to collect instantaneous circadian and cell cycle rhythms (Fig. 2H). The untreated cells exhibited an initial peak in circadian phase coherence after the phase resetting followed by a desynchronization (Fig. 2I). The treated cells reached the peak faster and started desynchronization at earlier time points, achieving the minimum phase coherence sooner (Fig. 2I). Hence, pharmacological inhibition of the TGF-β pathway accelerated the loss of circadian synchronization.

To evaluate the extent of intracellular communication between the circadian and cell cycle for untreated cells, we depicted their phases which revealed robust patterns of phase-locking (Fig. 2J). This observation implies that well-coupled circadian oscillators maintain a strong phase-locking with the cell cycle. However, in instances where the communication among circadian clocks is perturbed with the TGF-β pathway inhibitor drug or manipulation of seeding density, the circadian phases no longer exhibited a discernible preference for the timing of division events, destroying the phase-locking patterns (Fig. 2J-top and S2D). To measure the difference in phase-locking patterns, we employed a correlation coefficient between the two-dimensional distributions (see Methods). This analysis revealed a dose-dependent reduction in similarity to the untreated scenario in response to TGF-β pathway inhibitor with similar outcomes in the reduction of the seeding density (Fig. S2E).

Moreover, the temporal dynamics of circadian desynchronization resemble the decoherence of phase differences between the circadian and cell cycle, along with the detuning of their periods (Fig. S2F). By splitting the experiment into two stages—one characterized by high circadian phase coherence and the other by low phase coherence—we can alter the overall phase-locking patterns between the circadian rhythm and the cell cycle (Fig. 2J-bottom). Collectively, these results suggest the possibility of manipulating the global phase-locking dynamics between the circadian rhythm and the cell cycle through the modulation of extracellular circadian coupling. This manipulation can be achieved using a TGF-β pathway inhibitor drug, adjusting the seeding density, or by time decoherence.

### Extracellular circadian synchronization modulates cell growth

To delve deeper into the impact of circadian extracellular coupling on the cell cycle, we modulated the extracellular circadian coupling with an inhibitor of the TGF-β pathway (LY2109761), by decreasing the seeding density or by genetically disrupting the circadian clock and monitored populational growth (Fig. 3A). We measured growth in terms of population confluence while manipulating extracellular coupling with the LY2109761 drug which showed dose-dependent saturation levels and doubling times (Fig. 3B and Table S1). Notably, cells with well-coupled circadian clocks proliferated faster compared to cells with disrupted extracellular circadian communication. At the population level, the circadian signal of untreated cells revealed a more robust coupling among cells, manifesting a long-term sustained oscillation, as opposed to treated cells with the extracellular inhibitor (Fig. S3A).

**Figure 3:**
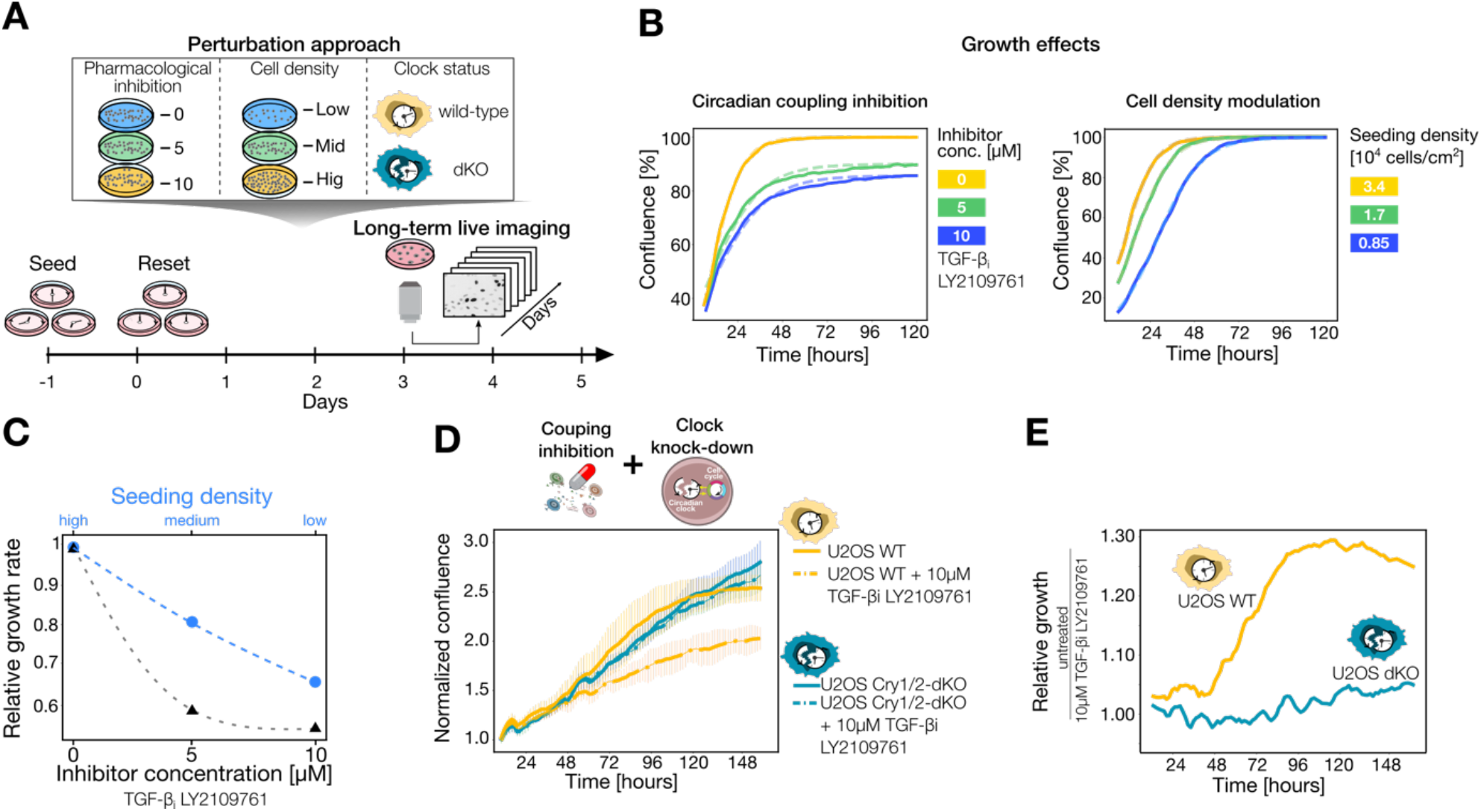
Extracellular circadian communication controls the growth rate. **(A)** Sketch of the experimental setup to disrupt the extracellular circadian coupling using the TGF-b pathway inhibitor LY2109761 (yellow: 0 μM, green: 5 μM, and blue 10 μM), by the seeding density (yellow: high density, green: medium density, and blue: low density) or genetical disruption of the circadian clock (yellow: wild-type, and turquoise: double-knockout of Cry1/2). **(B)** Left: growth measured in confluence for different concentrations of the extracellular circadian coupling inhibitor (yellow: 0 μM, green: 5 μM, and blue 10 μM). Right plot: growth measured in confluence for different seeding densities (yellow: 34000 cells/cm^2^, green: 17000 cells/cm^2^, and blue: 8500 cells/cm^2^). The curves were fitted (dashed lines) to a sigmoidal growth function to extract the growth rates (see Methods and Table S1). **(C)** Relative growth rates from Fig. 3B (for values and errors see Table S1). The dashed lines correspond to the fitting of the growth rates to an exponential function. **(D)** Normalized confluence of the U2OS cell lines: wild-type (yellow) and double-knockout of Cry1/2 (turquoise). Untreated cells are represented as a continuous line and treated cells with inhibitor LY21097621 are shown as dashed lines. The corresponding growth rates from the fitting to the sigmoidal growth function are shown in Table S2. **(E)** Relative growth measured by relative confluence for wild-type cells (yellow) and the double-knockout: (turquoise) between untreated and 10 μM LY2109761 treated conditions. The corresponding growth rates from the fitting of the sigmoidal growth function are shown in Table S4.

These results could be attributed to a circadian modulation of cellular growth or other potential effects of the TGF-β pathway inhibitor drug. To distinguish between these scenarios, we computed the growth measured by confluency while modulating the extracellular circadian coupling via the seeding density. We hypothesized that reducing the quantity of extracellular circadian signals in the media and, hence the extracellular circadian coupling, would lead to reduced growth. Cells with the lower seeding density demonstrated shorter doubling times, akin to the effect observed with the highest dose of LY2109761 (Fig. 3B). High-seeded cells exhibited a robust and prolonged circadian oscillation compared to the low-seeded which are associated with higher damping and lengthening of the circadian period (Fig. S3B).

To understand whether the observed growth differences might potentially be ascribed to circadian modulation we tested the LY2109761 effect in two U2OS cell lines: one wild-type with a well-functioning circadian clock (used in the aforementioned experiments) and a variant with a Cry1/2 double knockout^25^ (Fig. 3A). To do so, we compared the growth of untreated cells from both cell lines for 5 days, which revealed similar proliferation rates (Figs. 3D and S3C and Tables S2 and S3). Then, we compared the impact of the drug in both cell lines by computing the relative growth between the treated and untreated cells in each case. This uncovered a higher impact in the cells with a well-functioning circadian clock (Fig. 3E). This experiment showed that the U2OS wild-type cells with the TGF-β pathway inhibitor proliferated less compared to the untreated whereas the double-knockout cells exhibited similar proliferation rates irrespective of the presence of the drug (Fig. 3E and Table S4). Collectively, these findings underscore the influence of extracellular circadian synchronization and coupling on cellular growth.

### Growth curves show 24-hour oscillations

To discern the source of the growth differences and to avoid the averaging effect of population measures, we examined single-cell proliferation while disrupting extracellular circadian coupling using the TGF-β pathway inhibitor LY2109761. The cells displayed a heterogeneous proliferation pattern, spanning from no proliferation to high proliferation (Fig. 4A). To understand the impact of the inhibitor on the cell cycle, we computed the intermitotic times (IMT) of the tracked cells throughout the entire recording. The distribution of the intermitotic times shifted to higher values as the concentration of the inhibitor increased (Fig. 4B).

**Figure 4:**
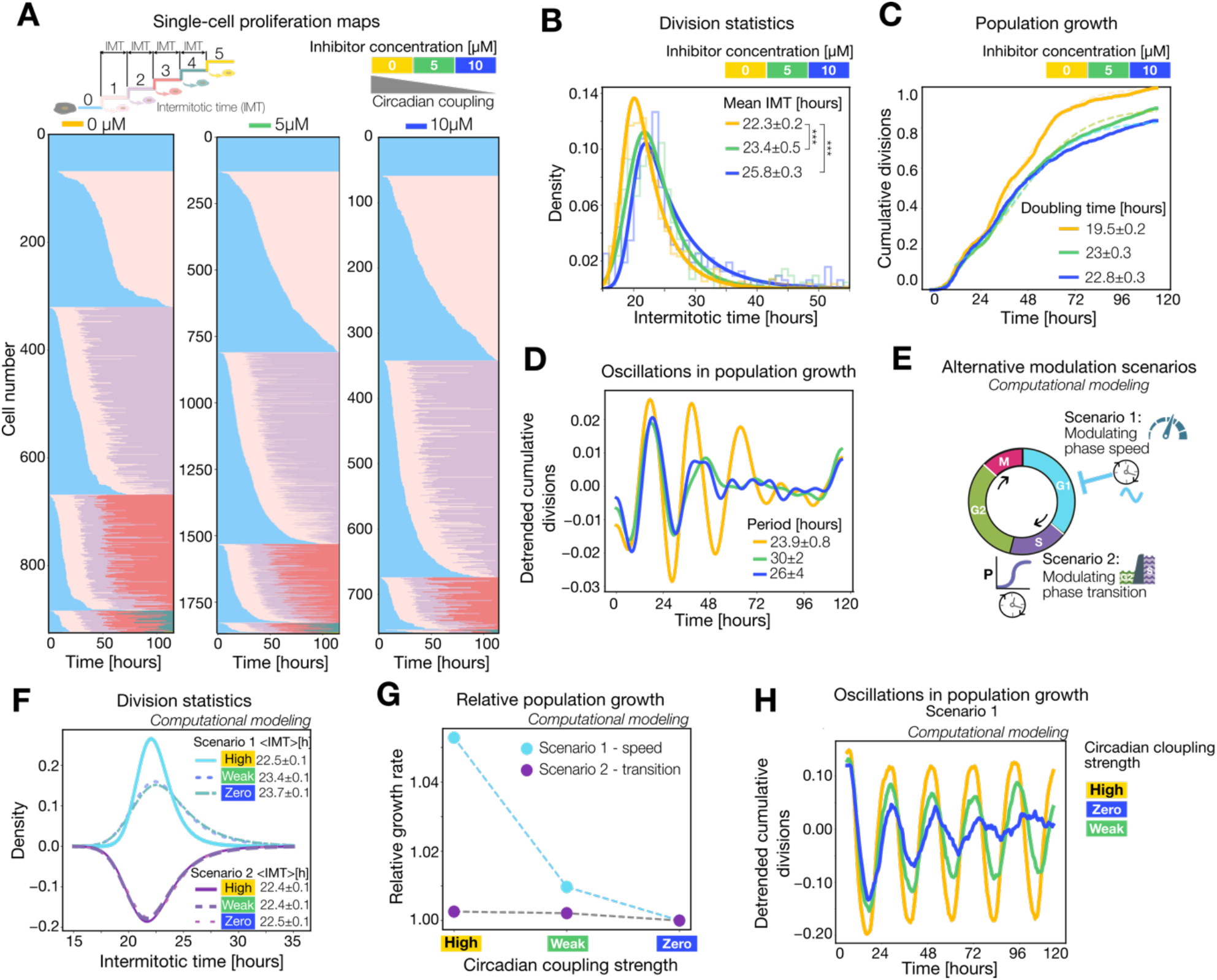
Growth curves exhibit 24-hour oscillations. **(A)** Proliferation plots of the single-cell level experiment from the same experimental design as Fig. 3A for different concentrations of the inhibitor LY2109761. Each row shows the division activity of an individual cell, with each mitotic event marked by a color transition (color code top left). Cells are clustered by their total number of divisions and then sorted by their first mitosis time. Sample sizes for 0 μM (n=923), 5 μM (n=1856), and 10 μM (n=759). **(B)** Distribution of intermitotic times of the cells that were present from the beginning of the recording for different concentrations of the inhibitor. The legend shows the mean with the corresponding error of fitting to an exponential modified Gaussian. Kolmogorov-Smirnov test was performed to compare the distributions. Sample sizes for 0 μM (n=654), 5 μM (n=910), and 10 μM (n=343). **(C)** Cumulative distribution of divisions for different concentrations of the inhibitor considering a bootstrapping to avoid sample size effects (n=750). The curves were fitted to a sigmoidal growth function (dashed lines). The legends show the doubling times corresponding to the growth rates shown in Table S5. **(D)** Detrended cumulative distribution of the division events from Fig. 4C for different concentrations of the inhibitor. The periods of the observed oscillations are shown in the legend. **(E)** Sketch of the growth mathematical model contemplating two possible scenarios: 1) modulating the phase speed and 2) modulating the phase transition. **(F)** Resulting fit to the histogram of intermitotic times. In blue color are shown the fits at different extracellular circadian coupling strengths, when the circadian rhythm is coupled to the G1 phase. In violet and on a flipped second axis are the distributions shown for coupling at the S transition. The legend shows the corresponding mean with the error from the fitting to an exponential modified Gaussian. **(G)** Relative growth rates of simulated cell cycles for different extracellular circadian coupling strengths, when the circadian rhythm is coupled to the G1 phase (blue) and the S transition (violet). Change is reported as the ratio between the growth rate of the circadian coupling strength and its value at zero strength. **(H)** Simulated detrended cumulative distribution of divisions for a well-coupled (yellow), mid-coupled (green), and uncoupled (blue) network of circadian clocks (left). The raw oscillations are shown in Fig. 4SF.

To study the proliferation evolution over time, we quantified the cumulative distribution of division events and observed a higher proliferation rate in the untreated cells compared to the treated (Fig. 4C and Table S5). The growth differences can be explained by longer intermitotic times and fewer divisions in the population of high concentrations of LY2109761, where the circadian clocks are weakly coupled. Strikingly, oscillations of approximately 24 hours were evident in the growth of the cell population with well-coupled circadian clocks, suggesting that cells proliferate synchronously with a periodicity of 24 hours (Figs. 4C-D). These 24-hour oscillations in growth exhibited quicker damping when the circadian coupling was reduced with the inhibitor drug, resulting in a longer period (Fig. 4D). Different seeding densities showed analogous outcomes (Figs. S4A-B and Table S6).

Next, to gain a better understanding of how extracellular circadian communication affects growth, we extend our mathematical model by incorporating a stochastic cell cycle model (see Methods). To account for the complex interactions between the circadian and the cell cycle, we test for two different scenarios: one with a coupling controlling the rate of progression and subsequently the length of the cell cycle, and another one with a coupling controlling the transition between phases and which can be seen as a gating mechanism. These coupling mechanisms can be included in both the G1 phase and S transition to model a cell cycle trace (Figs. 4E and S4C). Coupling in the G1 phase gives rise to results similar to the ones seen in the single-cell experiments (Figs. 4B-D), where intermitotic period distributions and growth rate are affected by an increasing drug concentration as a consequence of the decrease in the extracellular circadian coupling strength (Figs. 4F-G and S4D). Conversely, when the coupling is added to the S transition, no substantial impact on growth or intermitotic times is observed (Figs. 4F-G and S4D). This suggests that the main driver of the impact of the drug can be explained by the cell cycle length. In particular, cases with an inherently long circadian period will cause the growth rate to change giving rise to long cell cycles.

## Discussion

To synchronize with cyclic environmental conditions, organisms have evolved a molecular endogenous timing system of 24 hours, the circadian clock, which couples extracellularly to maintain coherence. Simultaneously, cellular division takes place to support growth and preserve homeostasis. Within numerous cells, these two internal rhythms operate concurrently with mechanisms of interactions yet to be uncovered. In this work, we address the impact of circadian synchronization on cell growth.

To understand the effect of extracellular circadian communication on the intracellular coordination between the circadian clock and cell cycle, we built a mathematical model using Poincaré oscillators. Our observations revealed that a well-coupled functional network of circadian clocks heightened the coordination between the cell cycle and the circadian clock. Furthermore, we found that the population phase-locking between the circadian clock and the cell cycle can be disrupted via extracellular circadian communication using a TGF-β secretory pathway inhibitor or the seeding density. Together, these findings highlight the potential role of the circadian clock on cell cycle regulation by showing that stronger communication between circadian clocks leads to a more robust phase-locking.

Further experiments indicated that by altering the circadian extracellular coupling with the TGF-β secretory pathway inhibitor, we can modify the growth rate of the population. To discard the attribution of growth differences to the toxic or proliferative effects of the inhibitor, we explored the seeding density effect as an alternative and observed analogous results. To gain a deeper understanding of the observed variations in growth, we investigated growth in two U2OS cell lines: a wild-type (U2OS WT) and a cell line with a disrupted circadian clock (U2OS Cry1/2-dKO BMAL1-Luc). The former cell line had a double knockout of two core clock genes, Cry1 and Cry2^26^. The addition of the TGF-β secretory pathway inhibitor changed the growth rate in the wild-type cell line. However, the drug produced no substantial changes in the double-knockout cell line. Hence, a population of cells with functional and synchronized circadian clocks exhibits enhanced proliferation compared to a cell line with a disrupted circadian clock. Taken together, these findings highlight the significance of extracellular circadian communication and synchronization in orchestrating growth. The cell cycle length will adapt to the circadian rhythm and will neither accelerate nor decelerate due to the 1:1 phase-locking between the circadian clock and the cell cycle, forcing proliferation rates to mirror the circadian periods.

On the population level, these growth differences could mask different cell cycle outcomes ranging from cell cycle arrest to death. To discern the proliferation behavior from other possibilities, we explored the division events at the single-cell level under the influence of the LY2109761 drug or by manipulating seeding density. In both instances of disrupting the circadian extracellular synchronization, we observed diminished proliferation rates compared to the untreated scenario with high seeding density. Moreover, the division events showed discernible oscillations indicating that cells divided synchronously and periodically every approximately 24 hours. These oscillations exhibited modified periods and quicker damping when disturbing the extracellular circadian communication.

By expanding the Poincaré model to include a stochastic cell cycle component, we gained insights into how circadian rhythms and population synchrony are interconnected. Specifically, we found that the length of the cell cycle, modulated by the coupled circadian rhythm, plays a key role in driving differences in growth rates for the model. When the circadian period became out of sync, so did the progression of the cell cycle, leading to a loss of synchrony in cell division across the population.

In summary, our study elucidates the interplay between the circadian clock and the cell cycle, and specifically, how the circadian clock exerts control over cellular growth. These initial findings lay the groundwork for unraveling the molecular mechanisms and further experiments are required to characterize the regulatory pathways involved. Potential contributors to this orchestration include Wee1, MYC, p16, and p21^20,27^. Inhibitors targeting these molecular mechanisms could be employed to evaluate their impact on the interplay between the circadian clock and the cell cycle. The growth and homeostasis of biological systems rely on the meticulous regulation of the cell cycle, which disruption leads to uncontrolled proliferation and, eventually, tumorigenesis. The potential ability to decelerate fast divisions within tumors to 24-hour cell cycle lengths using extracellular circadian clock communication holds significant promise for various cancer therapies.

## Acknowledgements

We would like to extend our gratitude to Huai-Yu Liou for his invaluable assistance in the establishment of the automatic tracking pipeline with ilastik. We thank the AMBIO imaging facility of the Charité Berlin for support in the acquisition of real-time fluorescence data. This work was financially supported by the German Federal Ministry for Education and Research (BMBF) through the e:Med Juniorverbund DeepLTNBC TP 3 - 01ZX1917C program. M.S.N. and M.H.J. acknowledge support from the Independent Research Fund Denmark (9040-00116B) and the Novo Nordisk Foundation (NNF20OC0064978). M.S.H. acknowledges support from the Lundbeck Foundation. C.E. thanks for the support from Berlin School of Integrative Oncology.

## Data availability

All the data generated and used for this investigation is available at the following link: https://itbfiles.biologie.hu-berlin.de/index.php/f/1427005.

The implemented code used for this study can be found here: https://github.com/Granada-Lab/Circadian-clock-cell-cycle.

## Ethics declaration: Competing interests

The authors declare no competing interests.

## Contributions

Conceptualization and investigation: NG, AEG

Methodology: NG, MSN, AEG

Experiments: MK, CE

Visualization: NG, AEG

Funding acquisition: AEG, MHJ, MSH

Project administration: AEG

Supervision: AEG

Writing – original draft: NG, MSN

Writing – review & editing: NG, MSN, CE, AEG

Intellectual support: UK, AK, MHJ, MSH, HH

## STAR Methods

### Materials and experimental setup

#### Cell culture

U2OS cells derived from ATCC HTB-96 were grown in RPMI medium containing l-glutamine (R8758 SIGMA), supplemented with 10% fetal bovine serum (FBS, Capricorn Scientific 12A) and 1% antibiotic–antimycotic solution (GIBCO 15240096).

#### Generation of reporter cell lines

details about the generation of the U-2 OS-*Cry1*/*Cry2*-dKO cell line can be found in the original publication^25^.

#### Fluorescent reporter cell lines

U2OS cells express a fusion protein NR1D1::VNP, comprising a nuclear localization signal and PEST element fused to the open reading frame of Venus, with the addition of the 116 N-terminal amino acids of NR1D1, as detailed in^26^. They were generously provided by M. Brunner from Heidelberg University. A singular subclone derived from this cell line was selected for subsequent experimental investigations.

#### Resetting

was achieved through incubation with 1 μM dexamethasone (Sigma dissolved in EtOH) for 30 minutes, employing standard tissue culture conditions. After the incubation period, cells underwent thorough washing with phosphate-buffered saline (PBS) in preparation for subsequent downstream applications.

#### TGF-β receptor inhibitor LY2109761

was prepared following the manufacturer’s guidelines, dissolved in the recommended solvents, and stored at −20°C. The drug treatment was administered at the specified concentrations during the seeding, after the resetting, and during imaging. The inhibitor persisted in the medium from their introduction throughout the entire duration of the experiment. The inhibitor LY2109761 and its solvent control were introduced into a conditioned medium (CM).

#### Microscopy

U2OS NR1D1::VNP reporter cells were cultured in FluoroBrite medium, composed of FluoroBrite DMEM (high glucose, without Hepes; Thermo Fisher Scientific #A1896701), supplemented with 2% FBS, 1% P/S, and l-glutamine (300 mg/liter; Thermo Fisher Scientific, #25030149) for long-term live-imaging.

Single-cell live imaging was achieved with high-resolution multichannel fluorescence microscopy using a high-resolution multichannel wide-field epi-fluorescence Nikon Ti2 microscope, equipped with an environmental chamber for cell culture (Okolab), and utilizing NIS-Elements visualization software (Nikon). Image acquisition occurred at 30-minute intervals, utilizing a 20× Plan Apo Ph2 DM λ magnification objective (Nikon), light-emitting diode illumination (Lumencor, SPECTRA X), and a scientific complementary metal-oxide semiconductor active-pixel sensor (sCMOS) PCO.edge camera (PCO), spanning multiple days.

The population live imaging was conducted through a prolonged, low-resolution incubator-embedded microscope (Incucyte, Essen Bioscience).

#### Image analysis

##### ilastik (v-1.4.0b21-OSX)^24^

First, we pre-processed the images to increase brightness and contrast through histogram equalization for each image sequence. To enhance the continuous visualization of the cell nuclei, we overlapped the images from the circadian and cell cycle channels. Then, we chose a representative image sequence to construct training files for segmentation and tracking in ilastik, following the approach outlined in^28^. The pixel classification workflow distinguished cells from the background and this segmentation output was employed in the tracking with learning workflow, utilizing optimized parameters tailored to our image dataset. For signal quantification, raw image sequences from the circadian and cell cycle channels were individually processed using ilastik as well. The object classification workflow was applied to extract mean signal intensities, and subsequent batch processing ensued.

##### Incucyte software (EssenBioScience, Incucyte 2022A)

In each frame, image analysis was conducted to count nuclei using the embedded Incucyte software within the Live-Cell Analysis Systems. This process facilitated the acquisition of confluency data as well as the quantification of reporter signal counts.

#### Computational methods

##### Filtering of signals

traces containing more than 100 frames (equivalent to 50 hours) were selectively retained for subsequent analyses. Discrepancies in nucleus size between consecutive frames were quantified to identify instances of erroneous tracking and merging cells. Traces exhibiting a nucleus size change surpassing 500 pixels were excluded, except in cases associated with cell division. Within each lineage, pairwise comparisons of circadian signal readouts were executed, leading to the removal of traces with a similarity exceeding 80%, attributed to late division events. Cells with at least one peak were kept for the subsequent data analysis. Only traces with continuous wavelet transform power exceeding 5% of the median power distribution were retained for subsequent analyses.

##### Division events

if the peaks in the gradient of the cell cycle signal exhibited abrupt changes and surpassed three times the mean of the gradient, they were defined as cell division events, highlighting discernible alterations in the Gemini fluorescent signal.

##### Preprocessing of single-cell signals

from the single-cell level experiment the individual circadian signals were extracted and detrended with the function sinc_detrend (with a cut-off period of 50h) and amplitude normalized with the function normalize_amplitude (with a window size of 50h) from the pyBOAT package of Python^29^. It uses a sinc filter that removes the periods larger than a certain cut-off. The cell cycle signals were not subjected to pre-processing. Finally, cells were ranked based on their circadian power, and only those exceeding a threshold of 10 were included in further analysis.

##### Continuous wavelet transform

we computed the continuous wavelet transform using the functions `compute_spectrum, get_maxRidge`, and `ridge_data` from the open-source software package `pyBOAT` to identify the predominant oscillatory elements characterized by the highest power through ridge detection for individual signals in a range of 16 to 40 hours.

##### Poincaré model

To simulate the circadian clock and cell cycle signals we employed Poincaré equations for the circadian clock with a mean field coupling (*extra*) and the cell cycle with a coupling (*intra*) to the circadian clock:

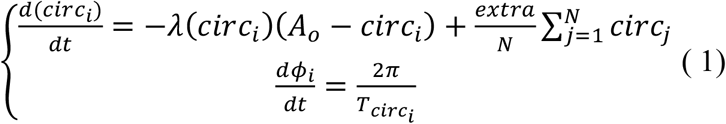

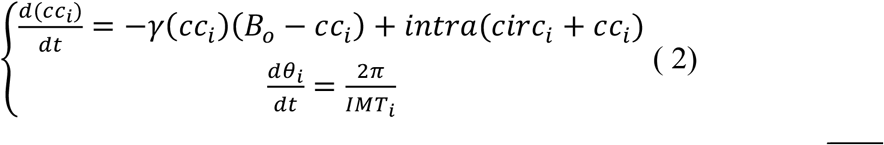

which were transformed to the Cartesian coordinates using 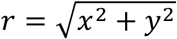 and 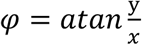 :

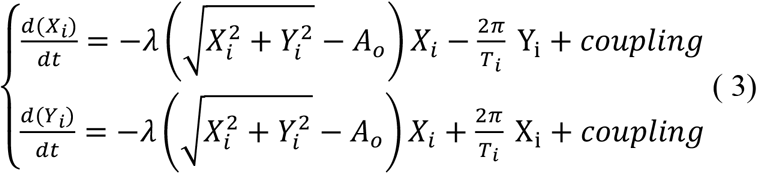

We defined the following parameters for the amplitude and relaxation of the circadian and cell cycle, respectively: *λ* = *A*_0_= *γ* = *B*_0_ = 1.

The periods were defined as *T*_*circ*_ *for* the circadian clock and *IMT as t*he intermitotic time for the cell cycle length. For Figs. 2F and S2A-C, the periods were obtained randomly from a Gaussian distribution with a mean of 22h (*T*_*circ*_, circadian clock) and 26h (*IMT*, cell cycle) and a standard deviation of 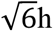 (both). For Fig. 2a, to simulate a network of circadian oscillators, we used Poincaré equations for the circadian clock with a mean field coupling (Eqs. 2 and 4).

The circadian clock periods (*T*_*circ*_) were obtained randomly from a Gaussian distribution with a mean of 24h and a changing dispersion between 0 and 10h. For Fig. 2b, a network of circadian oscillators was simulated using equations 2 and 4 with the periods obtained randomly from a Gaussian distribution with a mean of 22h and a dispersion of 6h. For Fig. 2E, we simulated a network of oscillators using equations 2-4. The periods were obtained randomly from a Gaussian distribution with a changing mean for the circadian clock (20-28h) and the cell cycle (28-20h) and a dispersion of 4h (both).

For Figs. 2A, 2F, and S2A-C, the initial conditions were randomly distributed between -1 and 1. For Figs. 2A and B, the initial conditions were randomly distributed between -1 and 1 and followed by an initial (first 10h) pulse of a Gaussian function of 0.01 amplitude, 0h mean, and standard deviation of 2h.

We used the following parameters for the different figures:

**Table 1.** Simulation parameters of the Poincaré model.

| **Figs.** | **$t_f$** | **$dt$** | **$N$** | ***extra*** | ***intra*** |
| --- | --- | --- | --- | --- | --- |
| **2A** | 120h | 0.1h | 20 | 0 - 0.04 | - |
| **2A-B** | 120h | 0.1h | 200 | 0 (blue), 0.0005 (green), 0.001 (yellow), and 0.05 (red) | - |
| **2G** | 300h | 0.1h | 200 | 0 – 0.05 | 0 – 0.25 |
| **2G** | 200h | 0.1h | 200 | 0 (left) and 0.05 (right) | 0 – 0.4 |
| **2F, S2A-B** | 120h | 0.1h | 200 | 0 (uncoupled), 0.0005 (weakly coupled) and 0.05 (coupled) | 0.1 |
| **S2C** | 120h | 0.1h | 200 | 0.05 (light pink and light purple) and 0 (dark-pink and dark-purple) | 0.1 (light pink and dark pink) and 0 (light purple and dark purple) |

##### Locking/coupling (Arnold) structures

For Fig. 2A, the synchronization index was measured with the following formula: 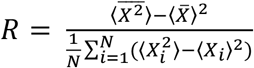

For Fig. 2G, an oscillator was considered to be locked internally when the circadian and the cell cycle had a stable phase difference after the transient stage, e.g. with a standard deviation smaller than 0.5 during the second half of the simulation. Then, we counted the fraction of how many oscillators were internally locked within the network and represented them with a heatmap.

#### Circadian properties

##### Fraction of oscillating cells

a cell was considered to be oscillating if it showed at least an oscillation of 24 hours with a mean wavelet power of 10 (a.u.). The peaks were found with the function find_peaks and the parameters (height=0, distance=32, prominence=0.1) from the preprocessed circadian signals.

##### Population circadian amplitude

The population oscillation represented in Figs. 2A, C, and D was obtained by summing the individual oscillations and dividing by N.

##### Population period

The instantaneous period of the population circadian oscillation was obtained using the continuous wavelet transform with the function compute_spectrum from the pyBOAT package of Python and smoothing the curve with the function sinc_smooth from the same package (with a cut-off period of 5h for the simulated periods or 10h for the experimental data).

##### Phase coherence

was computed with the following formula

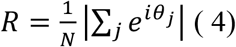

for Figs. 2B and S2 B, C and F, or it was plotted using the function get_ensemble_dynamics from the same pyBOAT package for Figs. 2G and S1E. This computational function employs the same formula as in (Eq. 5).

##### Single-cell periods

the instantaneous periods were extracted using the continuous wavelet transform from the preprocessed circadian signals. Then, the instantaneous periods were depicted in Figs. S1A and B using the function boxplot from the Python Seaborn package^30^.

#### Circadian and cell cycle locking metrics

##### Phase differences

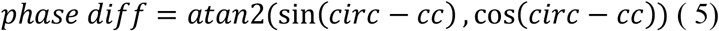

To compute the phase coherence of the phase differences, we used Equation 5 taking the phase differences using the above formula. For the mean value of the phase coherence of phase differences defined as the locking index, we avoided the transient stage and took the average from the second part of the simulation (Fig. S2A).

##### Phase space histograms

the phases were normalized between 0 and 1 by dividing by 2π. Then, a two-dimensional histogram was created with the NumPy^31^ function histogram2d and plotted with the imshow function from Matplotlib with a bilinear interpolation for simulations or by using the function hist2d from Matplotlib for experimental data.

##### Correlation coefficients between histograms

we established the untreated high-seeded cell population’s phase representation histogram as a reference. Then, we conducted a comparative analysis of alternative experimental conditions against this baseline by computing the correlation coefficient between flattened 2D histograms with the function corrcoef from NumPy. We flatted the 2D histogram by converting it into a 1D array while preserving the order of the elements with the flatten function in Python.

##### Normalized phase coherence and detuning

the circadian phase coherence was computed with equation 5 and normalized to the maximum value. The phase differences between the circadian and the cell cycle were computed using Equation 6. The phase coherence of these phase differences was computed using Equation 5 and normalized to the maximum value. The detuning between the circadian and the cell cycle was computed as the population average from the single-cell period difference between the circadian and the cell cycle.

#### Growth metrics

##### Growth curves

were measured by the cell confluence using the Incucyte software and were smoothed using the function savgol_filter (and the parameters: window_length=9, polyorder=2) from the Python package SciPy^32^ signal (Figs. 3B, E, and S3C).

##### Fitting growth/division curves

a sigmoidal growth function 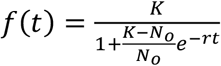 was used to fit the growth/cumulative division curves, where K denotes the growth saturation, *N*_*0*_ *is t*he initial growth and r is the growth rate. The fitting was achieved using the function curve_fit from SciPy signal library. From the growth rates we obtained the doubling times with the following formula 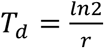 and the error was computed using the error propagation formula.

##### Proliferation plots

division matrices were constructed to encode division events (assigned a value of 1) and instances of no-proliferation (assigned a value of 0). Cells were grouped based on their total number of divisions, and within each division group, they were sorted according to the time at which the initial mitosis occurred (Fig. 4A).

##### Intermitotic distributions

the intermitotic times were obtained from the individual cell cycles using the continuous wavelet transform. Then, we chose the cells that were tracked from the beginning of the experiment for each condition and built a histogram using the Python function histplot from the Seaborn package.

##### Cumulative distribution of division events

a random equal sample of cells was bootstrapped from each condition to sum all the condensed division events of the cells over time to avoid potential sampling size effects. Then, we computed the accumulative sum of these division events with the Python function cumsum. This process was iterated 100 times to obtain the average cumulative distributions. Finally, all the conditions were normalized to the maximum of the cumulative distribution of the untreated condition.

Detrended cumulative distribution of division events and periods: the cumulative distribution of division events was detrended using the function sinc_detrend (with a cut-off period of 40h) and smoothed with the function sinc_smooth (with a cut-off period of 10h) from pyBOAT. The period was extracted with the continuous wavelet transform using the function compute_spectrum (from pyBOAT) in a range between 16 and 32 hours and presented in a boxplot.

##### Relative growth

was obtained by dividing the untreated confluence by the treated and smoothed with the function savgol_filter (and the parameters: window_length=9, polyorder=2) from the SciPy signal package of Python.

### Cell cycle mathematical model

To model the cell cycle, we wanted to keep it simple, yet being able to respond diversely depending on how the coupling between the cell cycle and a circadian rhythm is configured. To do so the cell cycle was broken down into two phases (G1 and G2), and two transitions between phases (S and M). When the circadian is coupled to one of the phases, it affects the speed of progression, while when coupled to the transition it modulates the probability for the cell to undergo a transition. Using a differential equation we can model this system as:

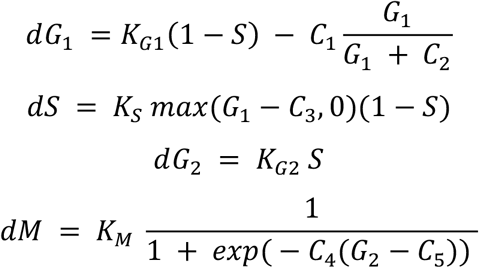

### Fitting

Using a Gillespie approach, we simulate the cell cycle as a stochastic system, where the randomness of the simulation can be changed by altering the “Volume” of the system, where a smaller volume will lead to a more stochastic result. Parameters were chosen for the system to have a self-sustained period of roughly 22h.

Circadian coupling can be introduced by multiplying one of the parameters (K_G1, K_S, K_G2 or K_M)

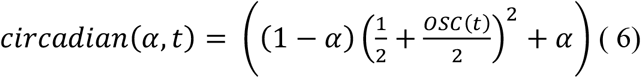

*α Is t*he coupling strength between the cell cycle and the circadian while OSC is the oscillating signal obtained from the Poincaré oscillators used previously (Eqs. 2-3).

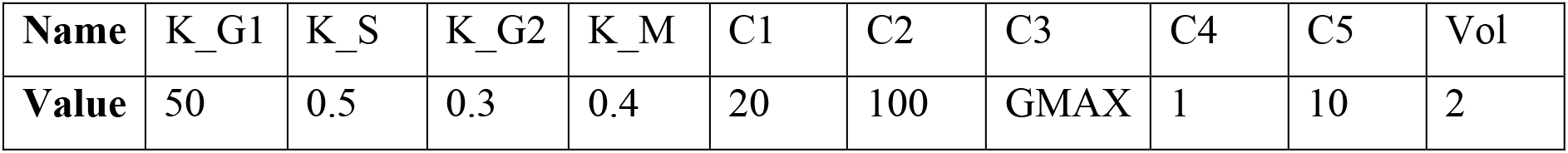

To initially match the circadian rhythm and cell cycle length, the parameter GMAX for each cell cycle was altered using

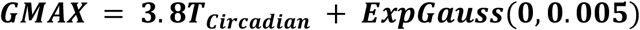

with ExpGauss being a random exponential modified Gaussian number drawn from Eq 8.

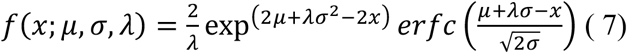

### Fit

Simulated cell cycles were obtained using 800 individually simulated cells, where each cell underwent cell division when passing the M transition in the cell cycle. The distribution (Fig 4f) of intermitotic periods was subsequently fitted to an exponentially modified Gaussian distribution (Eq. 8), by using a binned *χ*^*2*^ method.

From these parameters, we can calculate the mean of the distribution as

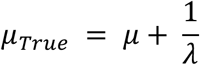

We likewise compute the error on the mean using standard error propagation using the parameter correlation found while fitting.

### Cumulative growth

Subdividing the time of simulation down into small time steps, we measure how many cells underwent mitosis during a given time step. Computing the cumulative number of divisions, we end up with a linear growth number indicating the total number of divisions of all cells at a given time. This signal is then detrended by subtracting a linear trend of the signal.

---

**Figure S1:**
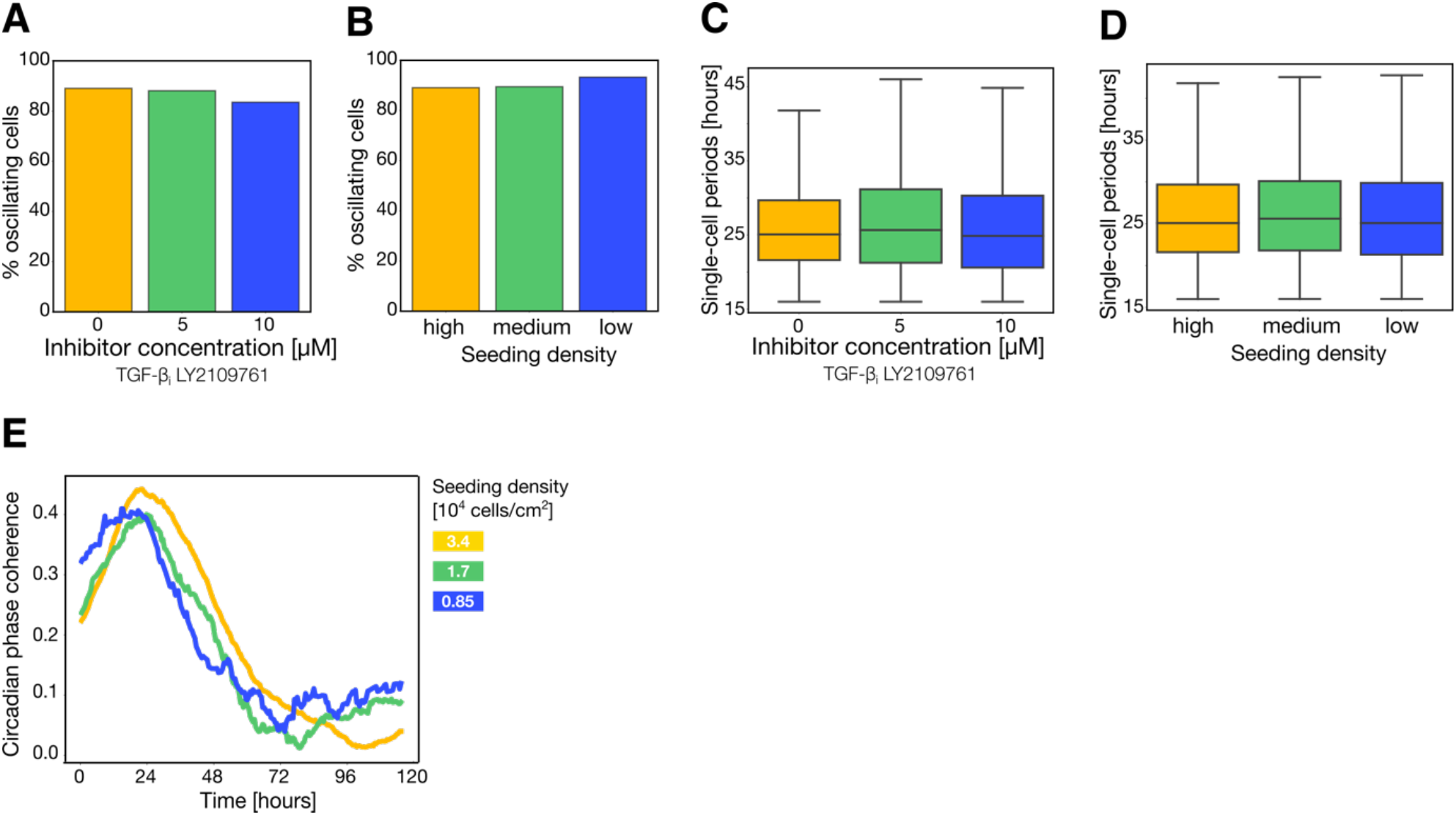
Circadian properties related to Figure 2. **(A)** Percentage of oscillating cells for different concentrations of the extracellular coupling inhibitor. **(B)** Percentage of oscillating cells for different seeding densities: high density (34000 cells/cm^2^), medium density (17000 cells/cm^2^), and low density (8500 cells/cm^2^ ). **(C)** Boxplot of the single-cell periods for different concentrations of the extracellular coupling inhibitor. **(D)** Boxplot of the single-cell periods for different seeding densities: high density (34000 cells/cm^2^), medium density (17000 cells/cm^2^), and low density (8500 cells/cm^2^ ). **(E)** Experimental circadian phase coherence for different seeding densities. Same sample size as for Fig. 2D.

**Figure S2:**
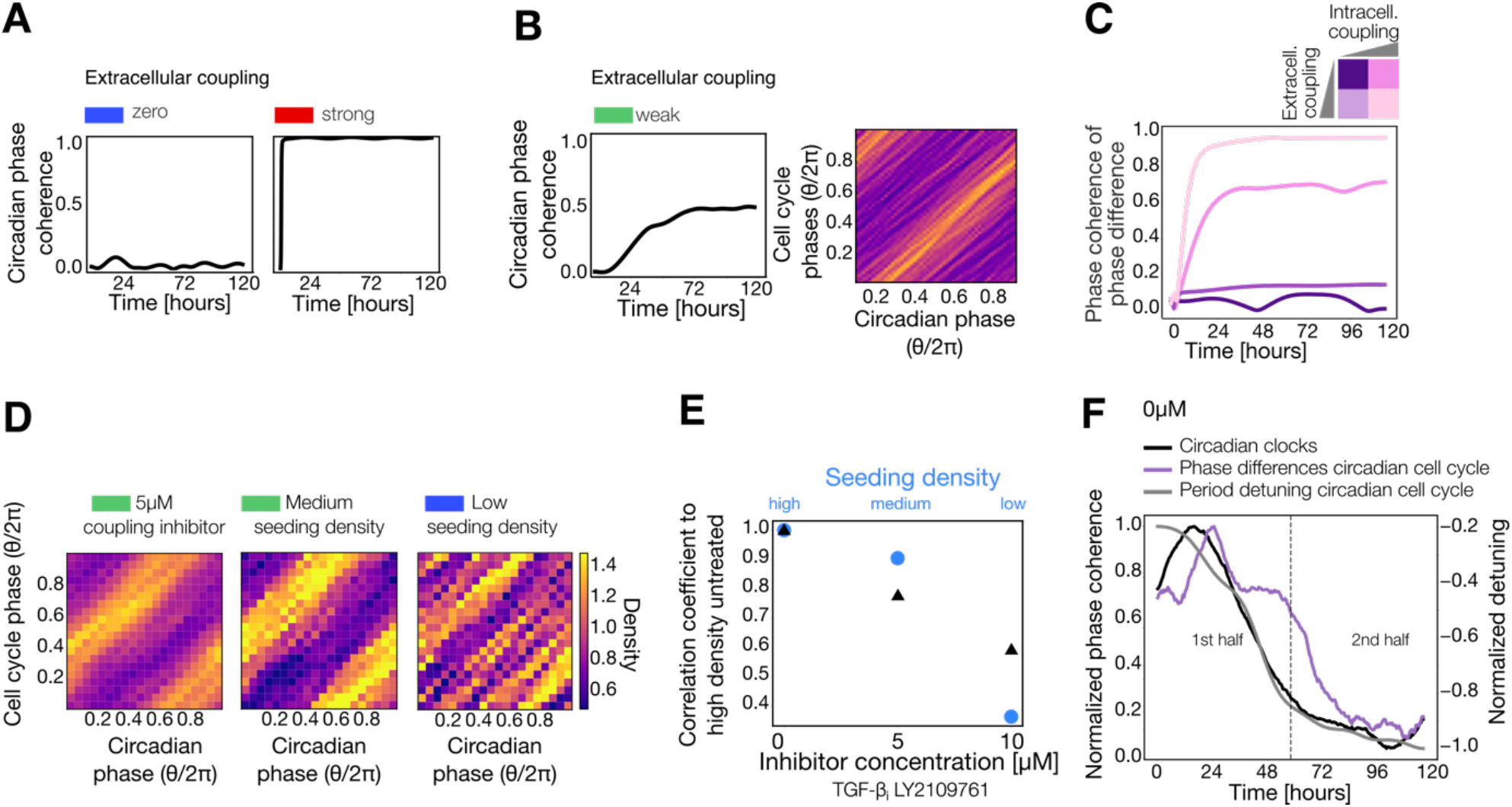
Circadian cell cycle locking characterization related to Figure 2. **(A)** Simulations of the circadian phase coherence of uncoupled and coupled circadian oscillators (see Methods). **(B)** Simulations of weakly coupled circadian oscillators with the corresponding phase coherence and the phase space representation of the circadian and cell cycle phases (see Methods). **(C)** Simulation of the phase coherence of the circadian and cell cycle phase differences of a network of oscillators with high extracellular and intracellular coupling (light pink), no extracellular coupling, and high intracellular coupling (pink), high extracellular coupling and no intracellular coupling (light purple) and no extracellular or intracellular coupling (dark purple). **(D)** Two-dimensional histograms of the circadian and cell cycle phases for 5 μM LY2109761 treated cells and medium- and low-seeded cells (17000 cells/cm^2^ and 8500 cells/cm^2^). **(E)** Correlation coefficients comparing the histograms of the circadian and cell cycle phases of the treated conditions with LY2109761 to the untreated (black triangles) and of the lower seeded densities (blue dots). **(F)** Normalized phase coherence of the circadian clocks (black) and the phase differences of the circadian and cell cycle (purple) to the maximum value of the untreated cells. The median period detuning of the circadian and the cell cycle (grey) was normalized to the minimum value. The dashed line splits the experiment into two identical halves.

**Figure S3:**
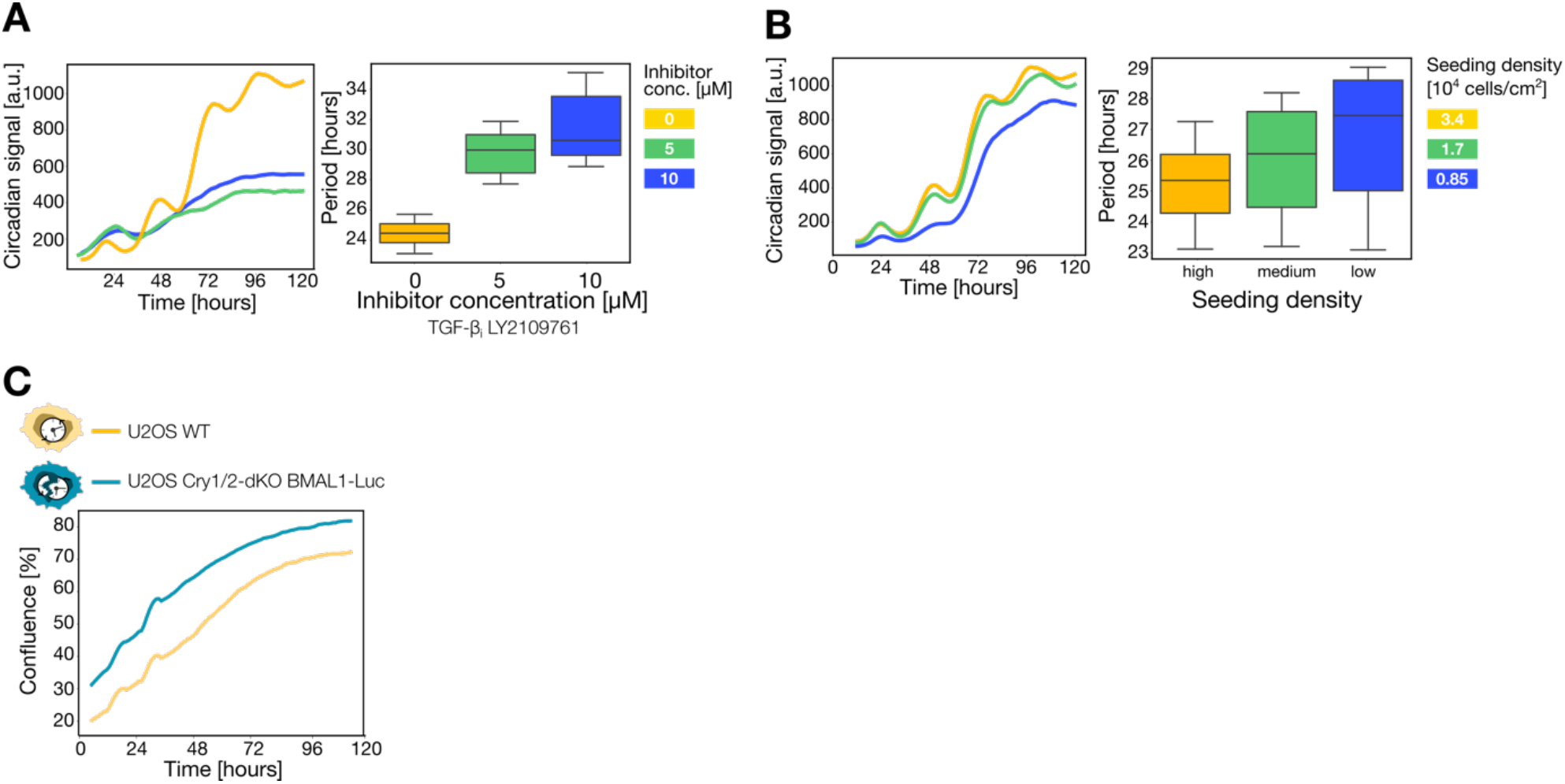
Population experiments related to Figure 3. **(A)** Right: circadian population signal from the experiment Fig. 3A for different concentrations of the inhibitor. Left: boxplot of the period of the population circadian signal for different concentrations of the inhibitor. **(B)** Left: circadian population signal from the experiment Fig. 3A for different seeding densities. Right plot: boxplot of the period of the population circadian signal for different seeding densities. **(C)** Growth measured by confluence for untreated cells of both U2OS cell lines (wilt-type: yellow and double-knockout: turquoise). The corresponding growth rates and errors obtained from the fitting to the sigmoidal growth function are shown in Table S3.

**Figure S4:**
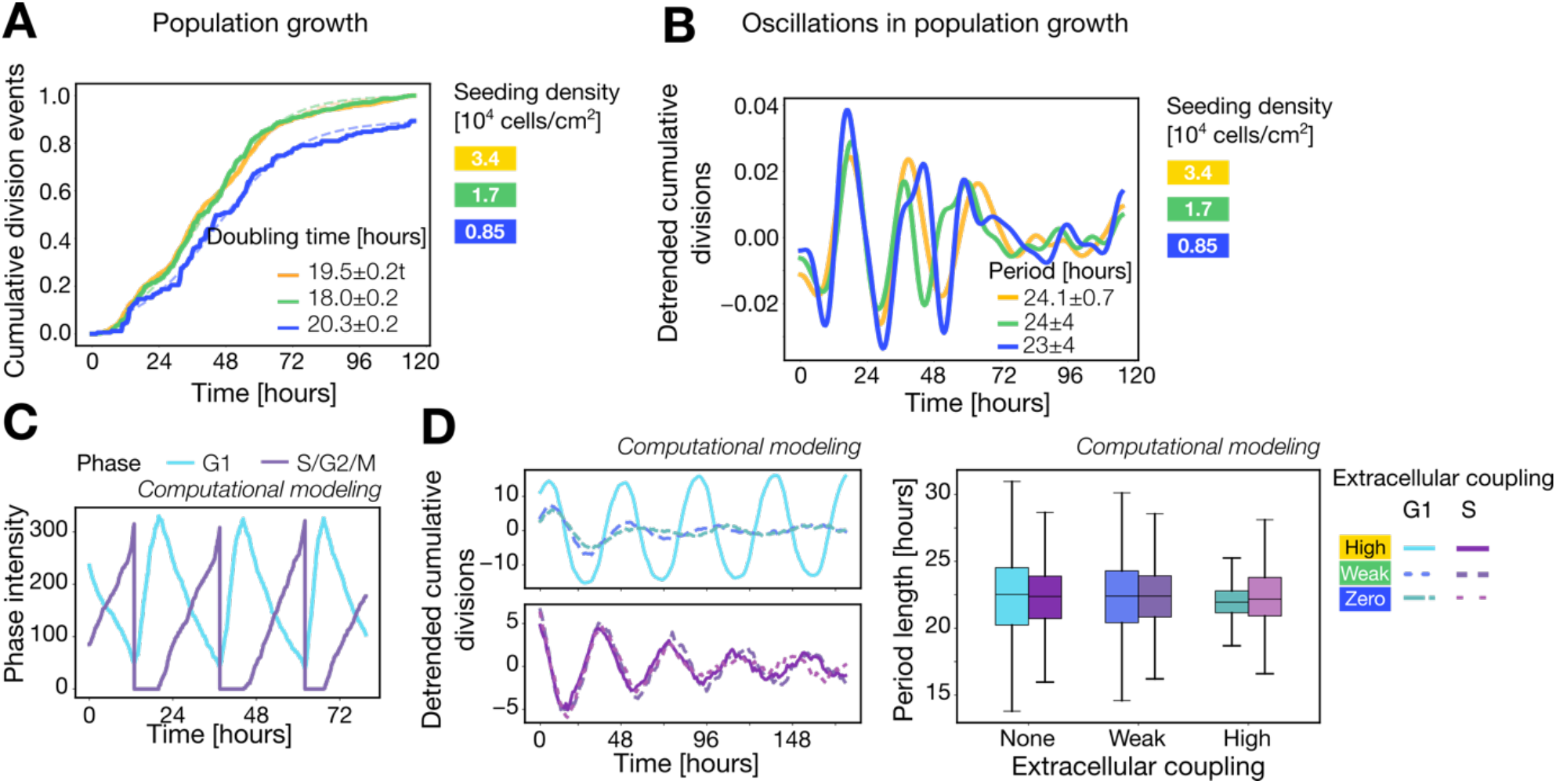
Additional plots to Figure 4. **(A)** Cumulative distribution of divisions for different seeding densities considering a bootstrapping to avoid sample size effects (n=89). The curves were fitted to the sigmoidal growth function (see Methods). The legends show the corresponding doubling times (for the growth rates see Table S4A). **(B)** Detrended cumulative distribution of the divisions for different seeding densities. The periods of the observed oscillations are shown in the legend. **(C)** Simulated single-cell traces corresponding to the G1 (blue) and S/G2/M (violet) phases. **(D)** Left: detrended cumulative divisions of multiple simulated cell cycles for different extracellular circadian coupling strengths, when the circadian rhythm is coupled to the G1 phase (blue) and the S transition (violet). Right: boxplot of the corresponding periods of the oscillations from the detrended cumulative divisions.

**Table S1:** Growth rates from the sigmoidal growth fitting corresponding to Fig. 3B.

| Condition<br>Seeding density | r [1/hours] | Condition<br>Coupling inhibitor | r [1/hours] |
| --- | --- | --- | --- |
| **Yellow**<br>**(high density)** | $0.127 \pm 0.001$ | **Yellow (0 $\mu\text{M}$)** | $0.127 \pm 0.001$ |
| **Green**<br>**(medium density)** | $0.105 \pm 0.001$ | **Green (5 $\mu\text{M}$)** | $0.074 \pm 0.001$ |
| **Blue**<br>**(low density)** | $0.0841 \pm 0.0004$ | **Blue (10 $\mu\text{M}$)** | $0.0841 \pm 0.0004$ |

**Table S2:** Growth rates from the sigmoidal growth fitting corresponding to Fig. 3D.

| Condition | r [1/hours] |
| --- | --- |
| **U2OS wild-type (0 $\mu\text{M}$)** | $0.023 \pm 0.001$ |
| **U2OS wild-type (10 $\mu\text{M}$)** | $0.0229 \pm 0.0004$ |
| **U2OS Cry1/2-dKO (0 $\mu\text{M}$)** | $0.028 \pm 0.001$ |
| **U2OS Cry1/2-dKO (10 $\mu\text{M}$)** | $0.0223 \pm 0.0004$ |

**Table S3:** Growth rates from the sigmoidal growth fitting corresponding to Fig. S3C.

| Condition | r [1/hours] |
| --- | --- |
| **U2OS wild-type** | $0.045 \pm 0.001$ |
| **U2OS Cry1/2-dKO** | $0.0449 \pm 0.0004$ |

**Table S4:** Growth rates from the sigmoidal growth fitting corresponding to Fig. 3E.

| Condition | r [1/hours] |
| --- | --- |
| **U2OS wild-type** | $0.023 \pm 0.001$ |
| **U2OS Cry1/2-dKO** | $0.001 \pm 0.001$ |

**Table S5:**
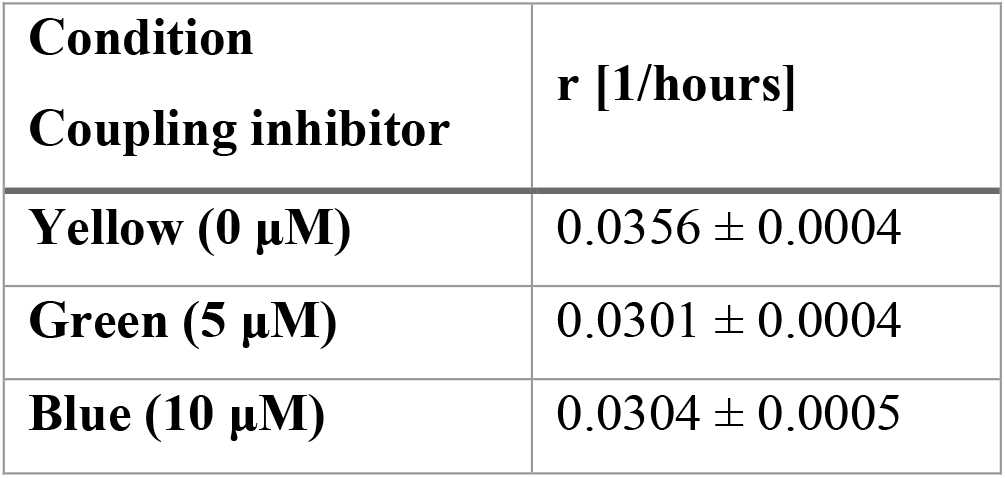
Growth rates from the sigmoidal growth fitting corresponding to Fig. 4C.

**Table S6:**
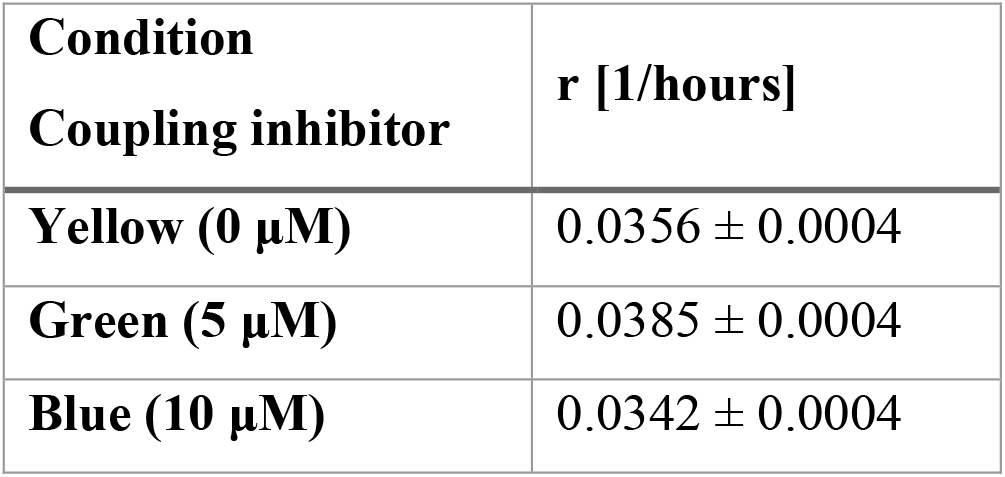
Growth rates from the sigmoidal growth fitting corresponding to Fig. S4A.

